# Physical interactions between pollen and pistil tissues mediate cryptic female choice in *Brassica rapa*

**DOI:** 10.64898/2026.08.04.742526

**Authors:** T. Chenin, E. Barbot, F. Rousset, A. Mignot, P. David, J. Tonnabel

## Abstract

Cryptic female choice - female-mediated bias in fertilization after mating - is well established in animals and can also occur in plants when multiple pollens compete on the same pistil. However, whether interactions between pollen and pistil tissues after pollen deposition contribute to this process remains unknown. Here, we experimentally test whether such interactions mediate cryptic female choice in the angiosperm *Brassica rapa*. We quantified fertilization success of pollen donors competing on the same pistil using paternity analyses, and in parallel, made semi-*in vivo* assays to measure pollen tubes trajectories emerging from the excised styles and growing toward unfertilized ovules for each donor-recipient pair. We show that pollen tube growth towards ovules predicts higher fertilization success under pollen competition. Thus, we document a previously unobserved mechanism of cryptic female choice based on physical interactions between male and female components of reproduction. In addition, different recipient plants favour different pollen donors, consistent with non-directional female choice. Plants with longer styles bias paternities more strongly towards the most successful pollen donor. Overall, our study demonstrates that interactions between pollen tubes and pistil tissues after pollen germination enable plants to bias paternity toward particular donors.

## Introduction

The study of sexual selection in plants has historically focused on traits influencing pollen export and pollen competition for ovule fertilization within pistil tissues (Tonnabel *et al*., 2026). This work has identified several sexually selected traits, including larger floral displays and floral or vegetative morphologies that enhance pollinator attraction or wind-mediated pollen dispersal during the pre-pollination phase (Parachnowitsch & Kessler, 2010; Karron & Mitchell, 2012; Tonnabel *et al*., 2019a; Brunet *et al*., 2021). Plant studies have converged with classical findings in animals (Janicke *et al*., 2016), by showing that male reproductive success generally increases with mate numbers, whereas this relationship is typically weaker in females (Johnson & Shaw, 2016; Tonnabel *et al*., 2019b; Kwok & Dorken, 2022; Barbot *et al*., 2023; Lavaut *et al*., 2023; Hou *et al*., 2024). Nonetheless, recent findings (Paschke *et al*., 2002; Barbot *et al*., 2023) suggest that females may also gain benefits from mating with multiple males in angiosperms, paralleling results in animals. Mating with several pollen donors provides an opportunity for sexual selection to occur after pollination. During the post-pollination phase, pollen traits such as pollen size or pollen tube growth rate typically influence gamete fertilization success (Snow & Spira, 1991; McCallum & Chang, 2016; Lankinen *et al*., 2017; Tonnabel *et al*., 2022). The post-pollination phase also provides females with the opportunity to exert cryptic female choice, a process by which female traits enable paternity bias toward particular males, which may confer genetic benefits to them. However, this process is well documented in animals but still rarely tested in plants (Pannell & Labouche, 2013).

Cryptic female choice refers to female-mediated bias in fertilization after mating and is typically associated with fitness benefits (Eberhard, 1996; Simmons & Kotiaho, 2007; Oliver & Evans, 2014; Rosengrave *et al*., 2016; Lüpold *et al*., 2016; Firman *et al*., 2017). In animals, it encompasses two non-exclusive processes favoring higher-quality partners based on variation in male traits, as predicted by Fisherian runaway or good-genes models (Fisher, 1915; Iwasa *et al*., 1991; Pomiankowski *et al*., 1991; Birkhead & Pizzari, 2002; Simmons & Kotiaho, 2007; Lüpold *et al*., 2016; Firman *et al*., 2017) or mates most compatible with the female’s own phenotype (Zeh & Zeh, 1997; Birkhead & Pizzari, 2002; Mays & Hill, 2004; Oliver & Evans, 2014; Rosengrave *et al*., 2016; Firman *et al*., 2017). Under the Fisherian runaway model, female choice evolves because alleles coding for female preference and preferred male traits become genetically correlated in the offspring of choosy females, thereby fueling a coevolution of both the preferred trait and preference (Fisher, 1915; Pomiankowski *et al*., 1991). A good-genes process may facilitate this coevolution, when the preferred male trait reflects overall genetic quality, allowing choosy females to produce fitter offspring in addition to attractive sons (Iwasa *et al*., 1991). A characteristic of these models is that females may be more or less choosy, but should all tend to favour the same males. Empirical evidence for post-copulatory female choice compatible with this pattern and mediated by variation in female reproductive tract morphology has been reported in animals, although direct evidence remains limited and largely restricted to insects (Miller & Pitnick, 2002; Simmons & Kotiaho, 2007; Lüpold *et al*., 2016; Firman *et al*., 2017). For example, larger spermatheca favor the paternity of males producing shorter sperm, a heritable trait associated with male fitness in *Onthophagus taurus* (Simmons & Kotiaho, 2007). Similarly, in plants, the first direct evidence of cryptic female choice (Chenin *et al*., 2026) showed that specific pistil morphology biased paternities toward pollen donors with specific pollen traits, although consequences for female reproductive success remain unknown.

A second form of cryptic female choice arises when the most successful male gamete donors vary among females (hereafter, compatibility-based female choice), whereby females favor fertilization by males whose phenotypes are most compatible with their own (Zeh & Zeh, 1997; Birkhead & Pizzari, 2002; Mays & Hill, 2004; Firman *et al*., 2017). This process may for example ensure inbreeding or outbreeding avoidance (Birkhead & Pizzari, 2002; Firman *et al*., 2017). In animals, evidence for compatibility-based female choice has been reported in internally and externally fertilizing species, where interactions between sperm and eggs, or associated female fluids can be directly observed (Gasparini & Pilastro, 2011; Oliver & Evans, 2014; Fitzpatrick & Evans, 2014; Firman & Simmons, 2015; Rosengrave *et al*., 2016; Gasparini *et al*., 2020). Fully factorial experimental designs in fishes and mussels have shown that ovarian fluids can differentially alter sperm velocity – a trait commonly linked to fertilization success – across specific male-female combinations, thereby generating significant male-by-female interaction effects (Urbach *et al*., 2005; Rosengrave *et al*., 2008, 2016; Evans *et al*., 2013; Oliver & Evans, 2014; Geßner *et al*., 2017; Poli *et al*., 2019). In internal fertilizers, several studies have reported greater fertilization success and both higher sperm velocity and viability in response to contact with ovarian fluids when males were unrelated rather than related to females in species such as guppies, mice and humans, findings compatible with compatibility-based female choice (Gasparini & Pilastro, 2011; Fitzpatrick & Evans, 2014; Firman & Simmons, 2015; Jokiniemi *et al*., 2020). More generally, female choice mediated by the Major Histocompatibility Complex (MHC), a widespread mechanism promoting reproduction between genetically dissimilar mates, has been shown to operate through post-mating processes (Løvlie *et al*., 2013; Firman *et al*., 2017; Lenz *et al*., 2018; but see Yeates *et al*., 2009 and Gasparini *et al*., 2015). An extreme form of male-by-female interaction occurs in many plants: the self-incompatibility systems, where pollen sharing alleles with the maternal plant is rejected by pistil tissues (Doucet *et al*., 2016; Guo *et al*., 2019; Genete *et al*., 2020). However, unlike cryptic female choice, self-incompatibility primarily functions as an all-or-nothing process that prevents self-fertilization or reproduction between kin. Even after rejection of pollen sharing self-compatibility alleles, plants may express cryptic preferences among the remaining pollen donors based on relative fitness benefits. A meta-analysis across 20 plant species revealed frequent deviations from random mating through elevated or reduced levels of inbreeding (Ismail & Kokko, 2019). Although such patterns may reflect homogamy for pre-mating traits such as phenology (Godineau *et al*., 2022), the potential contribution of pollen-pistil interactions remains unknown.

Post-pollination sexual selection in plants has been studied almost exclusively through the lens of pollen competition, despite growing evidence that this phase of reproduction can also enable female choice (Tonnabel *et al*., 2021, 2022; Chenin *et al*., 2026). Most angiosperms are highly polyandrous, and the number of pollen grains deposited on stigmas typically exceeds the number of ovules, creating competition among pollen donors for fertilization (Pannell & Labouche, 2013). Pollen tube growth within pistils involves a complex molecular dialogue that alters the gene expression in both male and female tissues (Tung *et al*., 2005; Qin *et al*., 2009; Chen *et al*., 2014). Pistils are not passive structures: they actively provide water, nutritive molecules, and guidance cues necessary for pollen tube growth and ovule targeting (Higashiyama *et al*., 1998; Higashiyama & Takeuchi, 2015; Johnson *et al*., 2019). Notably, pollen tubes that grow through pistil tissues locate ovules more effectively than pollen germinated exclusively *in vitro*, suggesting that pistil tissues capacitate pollen tubes to locate ovules (Higashiyama *et al*., 1998; Palanivelu & Preuss, 2006; Stewman *et al*., 2010). By modulating the production or composition of nutritive and guidance molecules, pistils may bias paternity towards some pollen donors, either in a common direction (the expected response under Fisherian or good-genes sexual selection) or in a female-specific direction (as expected from compatibility-based selection). Pistil morphology may further modulate these processes: longer pistils and larger stigmas may intensify pollen competition by increasing, respectively, the distance to reach ovules and the number of competing pollen grains (Tonnabel *et al*., 2021; Chenin *et al*., 2026). Although molecular studies have examined how pistils and pollen tubes interact, these processes have never been related to relative fertilization success of various pollen donors in competition, leaving a gap in our understanding of cryptic female choice in plants.

In this study, we measured variation among pollen donors in pollen tube growth and ovule-targeting ability following interactions with pistil tissues, and tested whether such variation underlies cryptic female choice in *B. rapa*. To do so, we created groups of five pollen donors and five recipient plants. On the one hand, we measured relative post-pollination fertilization success of the five donors on each of the five recipients by manually pollinating recipient stigmas with a mix of the five pollens, and then assessing paternities of the resulting seeds. On the other hand, we quantified for each donor-recipient pair separately, pollen tube growth and orientation toward unfertilized ovules after pollen tubes have traversed pistil tissues, using semi-*in vivo* assays (Figure 1; Higashiyama *et al*., 1998; Palanivelu & Preuss, 2006; Stewman *et al*., 2010; Lorbiecke, 2012). In these assays, pollen from a single donor was deposited onto a freshly excised pistil from a recipient plant and placed onto a growth medium (pollen and pistils were collected from flowers not used to assess paternity success). Pollen tubes first grew through stigmatic and stylar tissues before emerging onto the medium, where eight unfertilized ovules had been arranged around the excised style base (Figure 1; Higashiyama *et al*., 1998; Palanivelu & Preuss, 2006; Stewman *et al*., 2010; Lorbiecke, 2012). This experiment followed a fully factorial design (Lynch & Walsh, 1998), with one assay for each donor-recipient combination, yielding a total of 325 combinations. After a standardized growth period in the semi-*in vivo* assays, we measured three pollen tube traits: tube length (growth ability), tube straightness (efficiency of ovule targeting), and remaining distance to the nearest ovule or micropyle (reflecting both growth and ovule targeting ability). Hereafter, we refer to these traits as “pollen-pistil interaction traits” because they may capture components of male-female interactions that cannot be assessed using simple *in vitro* approaches on male and female organs separately. To test this hypothesis, we also measured *in vitro* pollen traits on donors and morphological pistil traits on recipient plants separately. We corrected in vitro traits and semi in-vitro interaction traits for various sources of experimental heterogeneity, and we call “scores” the inferred corrected trait values for each individual or donor-recipient pair.

**Figure 1.**
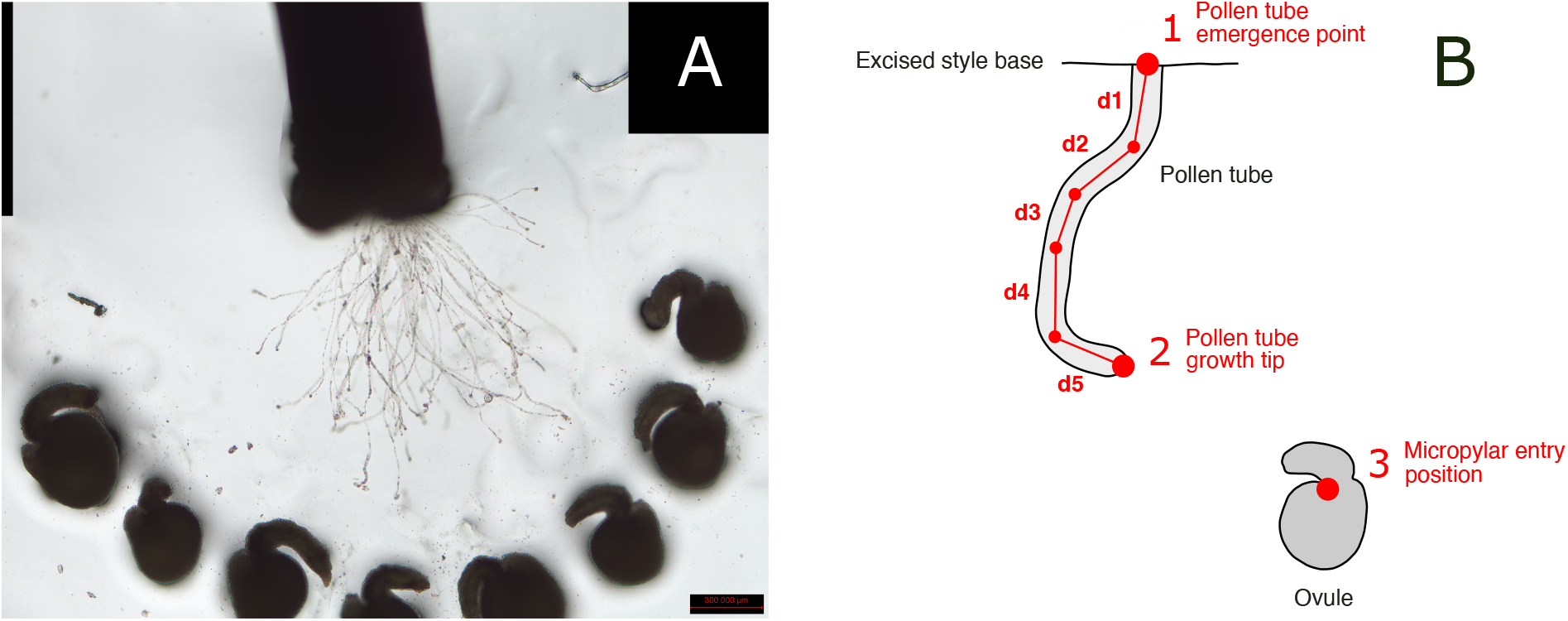
Microscope image of semi-*in vivo* assays and schematic representation of pollen tube growth toward an ovule. (**A**) Microscope image of pollen tubes emerging from an excised style base and growing in the medium (see Materials and Methods). The eight unfertilized ovules were manually arranged in a semi-circle around the excised style base. Five pollen tubes per image were randomly selected for pollen-pistil interaction score measurements. Scale bars: 300 µm. (**B**) Schematic representation of pollen tube growth and orientation toward an ovule. Pollen tube length was calculated as the sum of consecutive linear segments (here for example d1–d5) from the pollen tube emergence point at the excised style base [1] to the pollen tube growth tip [2]. Remaining distance to the nearest micropyle was defined as the Euclidean distance from the pollen tube growth tip [2] to the nearest micropyle [3]. Pollen tube straightness was calculated as Euclidean distance between the pollen tube emergence point [1] and the pollen tube growth tip [2] divided by pollen tube length.

Under cryptic female choice, we predicted that pollen donors whose pollen tubes grow farther, straighter, and closer to ovules after interaction with pistil tissues (stigma and style), would achieve higher fertilization success under pollen competition. Under a directional process, pollen-pistil interaction scores should be high or low for a given pollen donor regardless of recipient identity. Under a compatibility-based process, pollen-pistil interaction scores of each pollen donor should differ between recipients. We further tested whether cryptic female choice based on pollen-pistil interaction traits depended on pistil morphology such as stigma area, style length and ovary length scores. Accordingly, we examined whether: (1) pollen-pistil interaction traits vary mainly in relation to pollen donor identity, pollen recipient identity, or the donor-recipient pair, (2) pollen-pistil interaction traits are predicted by *in vitro* pollen and pistil scores previously demonstrated to underly cryptic female choice in *B. rapa* (Chenin *et al*., 2026), (3) pollen-pistil interaction scores predict paternal fertilization success, thereby testing their involvement in cryptic female choice processes, and (4) interact with pistil morphology to shape fertilization outcomes. Our results show that pollen tube growth following pollen-pistil interactions predicts fertilization success through donor-by-recipient interactions rather than intrinsic donor quality, and is further amplified by style length, consistent with compatibility-based cryptic female choice. The involved pollen-pistil interaction traits are not predicted by *in vitro* pollen and pistil scores and therefore mediate a distinct mechanism of female choice in *B. rapa* from the previously identified ones (Chenin *et al*., 2026).

## Results

### Pollen tube length after interaction with pistil tissues predicted fertilization success

We tested whether pollen-pistil interaction traits depended on pollen donor identity, recipient identity, or their interaction while correcting for various sources of experimental heterogeneity (Table 1).For both pollen tube length and the remaining distance traits, the interaction between donor and recipient was the major component of variance (9.63% and 36.2% respectively; Table 1). Pollen donor identity, recipient identity, and their interaction collectively explained 17.3% of the total observed variance in the remaining distance trait and 38.85% in pollen tube length score. In contrast, the pollen tube straightness trait showed only residual error and no detectable variance among individuals or donor-recipient pairs (Table 1). For pollen tube length after interaction with pistil tissues, in particular, variance associated with pollen donor identity was near zero (0.21%) and negligible in comparison to donor-recipient interaction variance (Table 1). After inferring pollen-pistil traits for each donor-recipient pair while correcting for experimental sources of variation, remaining distance to the nearest micropyle and tube straightness scores were negatively correlated with pollen tube length score (Figure S1).

**Table 1.**
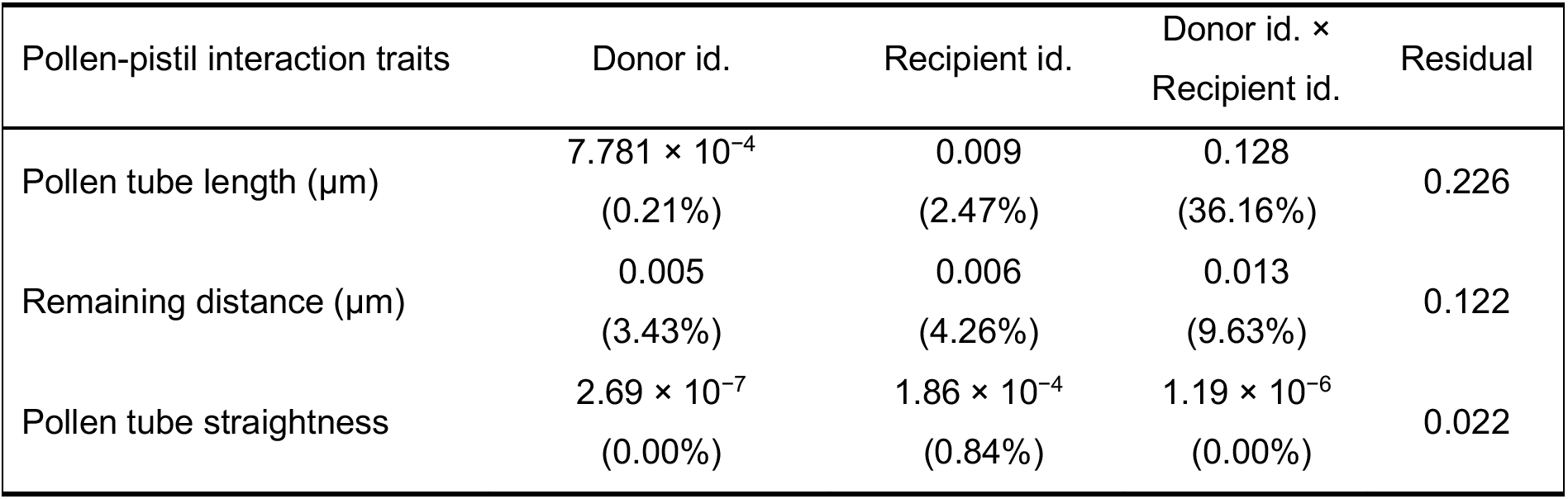
Variance in pollen-pistil interaction traits explained by pollen donor identity, pollen recipient identity, and their interaction. Pollen-pistil interaction traits were analyzed as response variables in separate generalized linear mixed-effects models (GLMMs) correcting for positional experimental sources of variation (see Table 3; see Materials and Methods for the random structure). Variance was further partitioned among pollen donor identity, pollen recipient identity, and their interaction included as random effects. For each pollen-pistil interaction trait, the percentages of observed variance explained by donor identity, recipient identity, and their interaction are reported in parentheses. Each percentage was calculated by dividing the amount of variance for each factor by the total amount of variance, defined as the sum of all variance components including residuals. Abbreviations. id.: identity.

Next, we tested whether pollen-pistil interaction scores influenced paternal fertilization success under pollen competition in a model with all three interaction scores as predictors. Higher pollen tube scores following pollen-pistil interactions were associated with higher fertilization success (slope β = 0.379, p < 0.001; Table 2; Figure 2A). By contrast, the remaining distance to the nearest micropyle and pollen tube straightness scores had no detectable effect on fertilization success (respectively: β = 0.010, p = 0.866, and β = 0.073, p = 0.310; Table 2; Figure 2B,C). In addition pollen tube length trait after interaction with pistil tissues was not predicted by *in vitro* pollen scores (pollen tube growth rate, germination rate, and pollen size; Table S1), recipient pistil scores (stigma area, style length, and ovary length; Table S1), or their interaction, although these scores have been previously identified as being involved in cryptic female choice in *B. rapa* (Chenin *et al*., 2026). Together, our results indicate that pollen tube length following physical interactions between pollen and pistil tissues does not mediate previously demonstrated effect of *in vitro* pollen scores, and therefore constitutes a new instance of cryptic female choice in *B. rapa*.

**Table 2.**
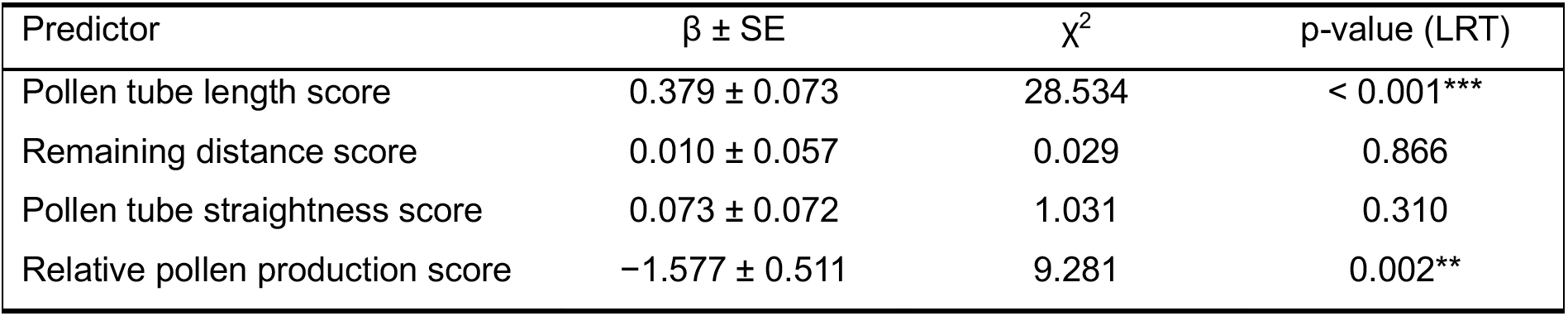
Estimated effects of pollen-pistil interaction scores and relative pollen production score on paternal fertilization success. Pollen-pistil interaction scores were measured from semi-*in vivo* assays for each pollen donor-recipient plant combination (*n* = 325). Pistils were excised at the style-ovary junction and placed on the medium before pollen application on the stigma. The model included the three pollen-pistil interaction scores as fixed effects, and the relative pollen production score of pollen donors as a covariate (see Materials and Methods). All variables were standardized before inclusion in the model. The significance of pollen-pistil interaction scores and relative pollen production score were tested using likelihood ratio tests (LRTs) comparing models with and without the effect. β values represent estimated effect sizes ± standard error. Degrees of freedom were equal to one for all model comparisons. Asterisks indicate statistically significant p-values: * p < 0.05, ** p < 0.01, *** p < 0.001. See also Figure 2.

| Predictor | $\beta \pm SE$ | $\chi^2$ | p-value (LRT) |
| --- | --- | --- | --- |
| Pollen tube length score | $0.379 \pm 0.073$ | 28.534 | < 0.001*** |
| Remaining distance score | $0.010 \pm 0.057$ | 0.029 | 0.866 |
| Pollen tube straightness score | $0.073 \pm 0.072$ | 1.031 | 0.310 |
| Relative pollen production score | $-1.577 \pm 0.511$ | 9.281 | 0.002** |

**Figure 2.**
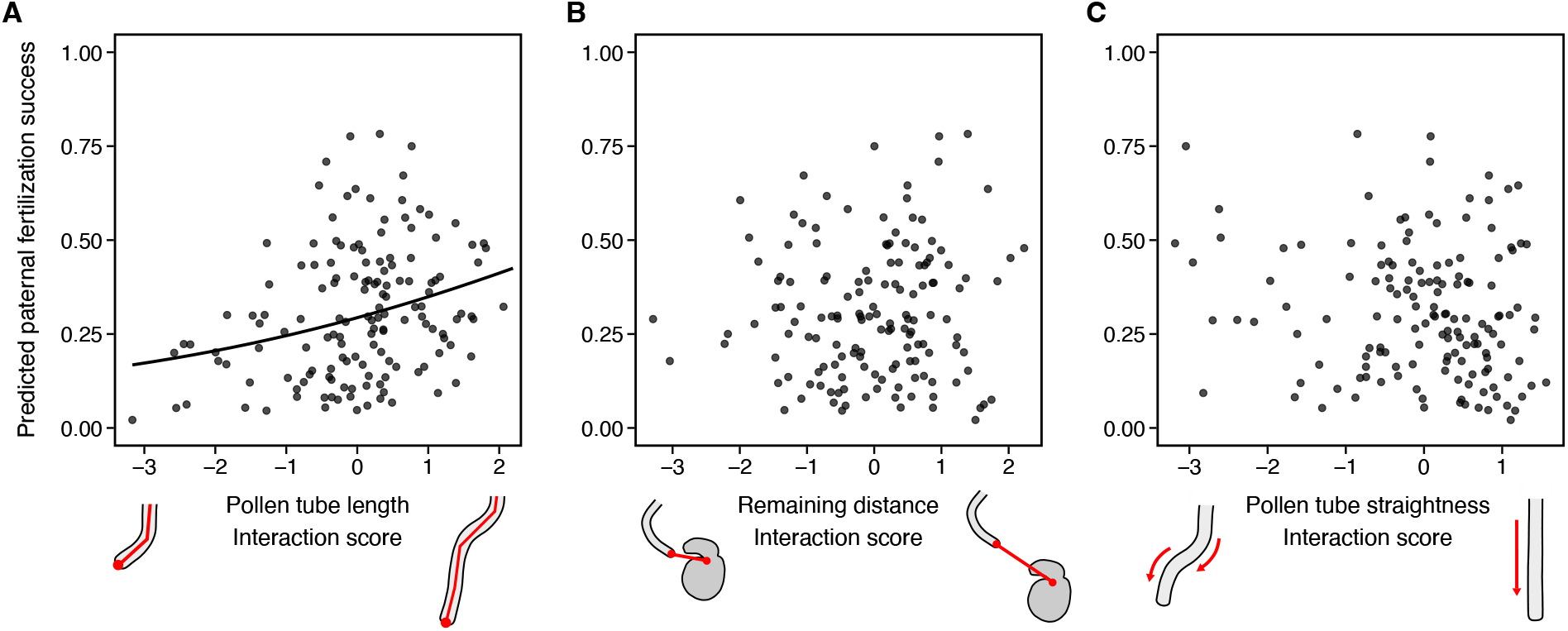
Pollen tube length score following pollen-pistil interactions predicts paternal fertilization success. Effects of the three pollen-pistil interaction scores on paternal fertilization success are shown for **(A)** pollen tube length, **(B)** the remaining distance to the nearest micropyle, and (**C)** pollen tube straightness after pollen tubes had traversed the stigma and style on paternal fertilization success. The model included all three pollen-pistil interaction scores as fixed effects along with relative pollen production score for each pollen donor as a covariate (see Materials and Methods). Pollen-pistil interaction scores were measured under our semi-in vivo protocol for each donor-recipient plant combination within the same mating group (n = 325). Pollen-pistil interaction scores correspond to centered and standardized latent values inferred from the semi-in vivo analysis (see Materials and Methods). Black dots represent the predicted fertilization success of a given pollen donor plant among five donors in competition on the same recipient (prediction from multinomial models). The black line shows model predictions for the significant effect of pollen tube length interaction score on paternal fertilization success **(A).** See also Table 2.

To ensure that the relationship between pollen-pistil interaction scores and fertilization success was not driven by potential confounding effects, we accounted for uncontrolled sources of variation in our statistical framework (see Materials and Methods). We first corrected pollen-pistil interaction traits for uncontrolled variation in the positional arrangement of pollen tubes and ovules in our semi-*in vivo* assays, effectively creating pollen-pistil interaction scores as mentioned above (Table 3; Materials and Methods). Pollen tube length and the remaining distance to the nearest micropyle increased when ovules were positioned farther from the excised style base (respectively: β = 0.088, p = 0.024, and β = 0.260, p < 0.001; Table 3), and pollen tubes emerging farther from the style center exhibited shorter length and longer remaining distances to the nearest micropyle (respectively: β = -0.051, p = 0.018, and β = 0.046, p = 0.001; Table 3). The total distance traveled by pollen tubes through pistil tissues before emerging on the growth medium (sum of stigma and style length) had no detectable effect on pollen-pistil interaction traits (Table 3). Furthermore, the effects of pollen-pistil interaction scores on paternal fertilization success were independent of variation in relative pollen production scores among pollen donors; we accounted for this factor as a covariate in all fertilization success models, because we did not fully control for the quantity of pollen deposited on recipient plants during manual pollinations (Table 2; see Materials and Methods). We observed a negative effect of relative pollen production scores on paternal fertilization success (β = -1.577, p = 0.002; Table 2). By accounting for uncontrolled variation in positional arrangement effects and pollen production, our analyses characterize the unbiased effects of pollen-pistil interaction scores on fertilization success.

**Table 3.** Estimated positional effects on pollen-pistil interaction traits. Pollen-pistil interaction traits were analyzed as response variables in separate GLMMs (see Materials and Methods). Three positional factors were included as fixed effects, corresponding to standardized mean distances between: (i) pollen tube emergence at the excised style base and the eight micropyles (mean distance to eight micropyles), (ii) pollen tube emergence at the excised style base and the center of the excised style base (distance to the center of the excised style base), and (iii) the sum of stigma and style lengths on the medium (pollen tube travel distance through pistil tissues). Random effects included pollen donor identity, pollen recipient identity, and their interaction (see Table 2; see Materials and Methods for the complete random structure). The significance of each positional experimental factor was tested using LRTs comparing models with and without the effect. β values represent estimated effect sizes ± standard error. Degrees of freedom were equal to one for all model comparisons. Asterisks indicate statistically significant p-values: * p < 0.05, ** p < 0.01, *** p < 0.001.

| Positional factors | $\beta \pm SE$ | $\chi^2$ | p-value (LRT) |
| --- | --- | --- | --- |
| **Pollen tube length ($\mu\text{m}$)** |  |  |  |
| Mean distance to eight micropyles | $0.088 \pm 0.036$ | 5.106 | 0.024* |
| Distance to the center of the excised style base | $-0.051 \pm 0.021$ | 5.637 | 0.018* |
| Pollen tube travel distance through pistil tissues | $-0.061 \pm 0.042$ | 1.934 | 0.164 |
| **Remaining distance ($\mu\text{m}$)** |  |  |  |
| Mean distance to eight micropyles | $0.260 \pm 0.019$ | 119.156 | < 0.001*** |
| Distance to the center of the excised style base | $0.046 \pm 0.014$ | 10.145 | 0.001*** |
| Pollen tube travel distance through pistil tissues | $-0.015 \pm 0.022$ | 0.415 | 0.520 |
| **Pollen tube straightness** |  |  |  |
| Mean distance to eight micropyles | $-0.002 \pm 0.006$ | 0.110 | 0.740 |
| Distance to the center of the excised style base | $0.007 \pm 0.006$ | 1.630 | 0.202 |
| Pollen tube travel distance through pistil tissues | $0.003 \pm 0.007$ | 0.174 | 0.677 |

### Style length modulates the fertilization advantage of most successful pollen donors

We tested whether the effect of pollen tube length score after interaction with pistil tissues on fertilization success depended on three uncorrelated characteristics of the recipient plant: stigma area, style length, and ovary length scores (Table 4; Figure S2; see Materials and Methods). Among these, only the interaction between style length score and pollen tube length score was significant (β_style length score x pollen tube length score_ = 0.208, p = 0.021; Table 4). As shown in Figure 3, pollen donors with higher pollen tube length scores achieved greater fertilization success in plants with higher style length scores, whereas differences among pollen donors were smaller in plants with lower style scores (Figure 3). These results indicate that longer styles reinforce the fertilization advantage of pollen donors with higher pollen tube length scores.

**Table 4.**
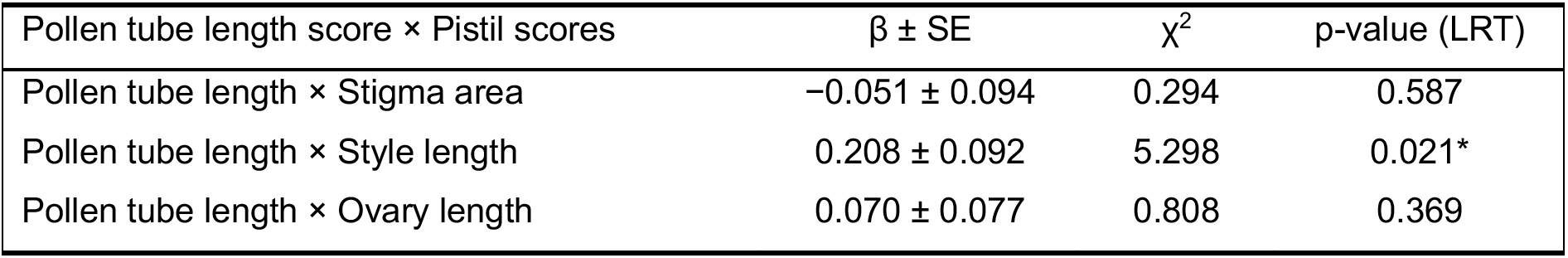
Estimated interactions between pollen tube length interaction score and stigma area, style length and ovary length scores. The model included pollen tube length interaction score as a fixed effect, along with its interaction with stigma area, style length, and ovary length scores, three uncorrelated components of pistil morphology (Figure S2; see Materials and Methods). Relative pollen production score for each pollen donor was included as a covariate in the model (see Materials and Methods). Pollen tube length interaction score was obtained for each pollen donor-recipient plant combination (semi-*in vivo* assays, *n* = 325), and pistil scores were obtained from several flowers of the recipient plant (*n* = 70). All variables were standardized before inclusion in the model. The significance of each interaction between pistil scores and pollen-pistil interaction scores was tested by comparing models with and without the effect using LRTs. β values represent estimated effect sizes ± standard error. Degrees of freedom were equal to one for all comparisons. Asterisks indicate statistically significant p-values: * p < 0.05, ** p < 0.01, *** p < 0.001. See also Figure 3.

**Figure 3.**
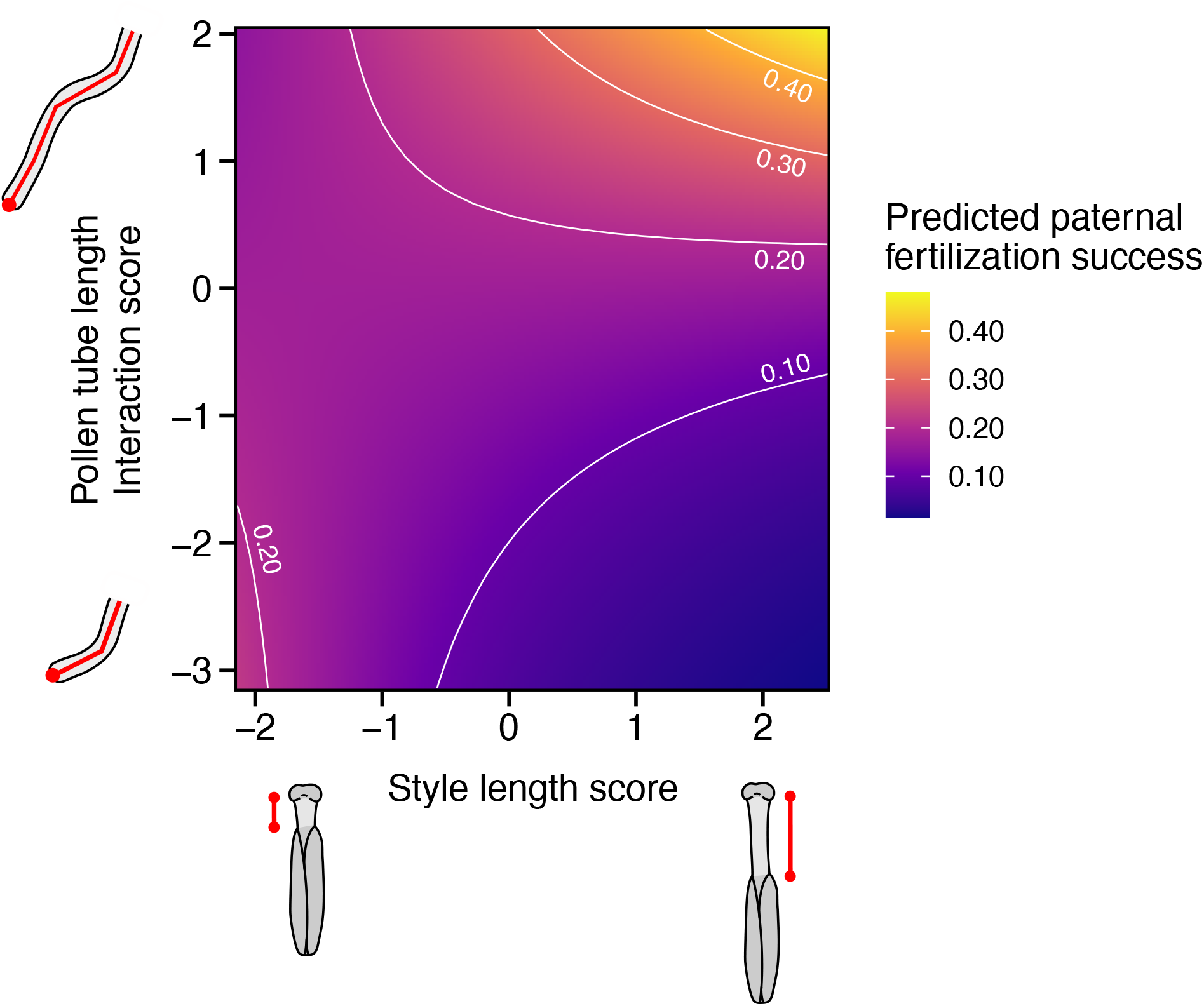
Style length score modulates the effect of the pollen tube length interaction score on paternity. Effect of the statistical interaction between pollen tube length interaction score (y-axis) and style length score (x-axis) on paternal fertilization success (color scale). The model included pollen tube length interaction score as a fixed effect, along with its interaction with stigma area, style length, and ovary length scores, three uncorrelated components of pistil morphology (Figure S2; see Materials and Methods). Relative pollen production score for each pollen donor was included as a covariate in the model (see Materials and Methods). We represented the only statistically significant interaction of the model, between pollen tube length interaction score and style length score, while holding all other fixed effects (none of which were significant) at zero, corresponding to their standardized mean values. Colors indicate predicted paternal fertilization probabilities, with yellow indicating higher predicted probabilities and purple lower probabilities. White contour lines delineate probability levels at 0.10 increments.

Our results regarding both the effect of pollen-pistil interaction scores and their interaction with pistil scores on fertilization success were robust to the exclusion of donor-recipient combinations in which the donor fertilized less than five percent of the recipient’s seeds (representing 34 donor-recipient combinations out of 174 for which we have pollen-pistil interaction trait measurements = 19.54%; Table S3–S4). This finding suggests that our results are not solely driven by the complete exclusion of certain pollen donors by recipient plants, as would be expected under self-incompatibility systems. Further, we found no evidence that inbreeding affected fertilization success, when inbreeding was measured by Loiselle’s kinship coefficient between donor and recipient, estimated from eight microsatellite loci (β = - 0.152, p = 0.748; see Materials and Methods).

## Discussion

We provide empirical evidence that the length of pollen tubes after traversing the stigma and style is positively correlated with fertilization success. This pollen-pistil interaction score does not reflect an intrinsic property of either the pollen donor or the recipient but instead emerges from their interaction. Moreover, it is not predicted by independently measured male and female scores, such as pistil or pollen morphology or pollen growth dynamics measured *in vitro*. Our results therefore complement recent empirical evidence that these independently measured pollen and pistil traits are involved in cryptic female choice in *B. rapa* (Chenin *et al*., 2026), thus suggesting that fertilization success is influenced both by individual reproductive traits and by emergent properties of pollen-pistil interactions. Our study advances this earlier work by providing an integrative assessment of the ovule-targeting abilities of pollen tubes following their development through pistil tissues, and illustrates how this physical interaction – an inherent component of plant reproduction – affects pollen competitive ability. These results therefore highlight a methodological limitation of approaches measuring male and female traits separately, as cryptic female choice processes are not fully captured by either *in vitro* pollen growth assays or morphological measurements alone. Moreover, we identify style length as a key pistil trait modulating the magnitude of cryptic female choice, reinforcing the importance of pistil morphology for post-pollination sexual selection in plants. Overall, our results suggest that cryptic female choice processes are more diverse than previously recognized, converging with findings from molecular biologists about the active role of pistil tissues during plant reproduction (Higashiyama *et al*., 1998; Higashiyama & Takeuchi, 2015; Johnson *et al*., 2019).

### Evidence for compatibility-based cryptic female choice mechanism

We show that variation in pollen tube growth following pollen-pistil interactions depended on which recipient plant a given pollen donor interacted with and that this pollen-pistil interaction influenced fertilization success under pollen competition. In the animal literature, the combination of these two results is generally interpreted as evidence that male gamete recipients differ in their preferences (Gasparini & Pilastro, 2011; Oliver & Evans, 2014; Rosengrave *et al*., 2016; Firman *et al*., 2017). These studies are consistent with so-called compatibility-based cryptic female choice (Zeh & Zeh, 1997; Birkhead & Pizzari, 2002; Firman *et al*., 2017), and our study documents the same type of process in plants. This pattern contrasts with assumptions of classical Fisherian or good-genes models of sexual selection, in which females are assumed to bias paternity directionally toward the same preferred males (Fisher, 1915; Iwasa *et al*., 1991; Pomiankowski *et al*., 1991). Our results parallel experimental studies in animals that have provided evidence for cryptic female choice by artificially recreating interactions between male gametes and female reproductive fluids (Gasparini & Pilastro, 2011; Oliver & Evans, 2014; Rosengrave *et al*., 2016; Alonzo *et al*., 2016). Using a comparable experimental approach, we obtained evidence that pollen-pistil interactions can favor specific combinations of compatible partners. These findings reveal a previously unrecognized source of paternity bias in plants and further support the view that pistil tissues are active participants in plant reproduction rather than passive structures (Moore & Pannell, 2011; Johnson *et al*., 2019; Tonnabel *et al*., 2021; Chenin *et al*., 2026).

The pattern of compatibility-based cryptic female choice observed here for *B. rapa* was not the mere consequence of the sporophytic self-incompatibility systems characteristic of the Brassica genus (Takayama & Isogai, 2005; Nasrallah, 2017). First, our interaction traits were measured after the pollen tube had emerged from the base of the style, whereas pollen grains sharing self-incompatibility alleles with female tissues typically fail to germinate or are arrested before reaching that stage (Nasrallah, 2019). Second, our results remained qualitatively robust after excluding donor-recipient combinations in which the donor sired fewer than 5% of the recipient’s seeds, suggesting that the observed pattern reflects quantitative differences in fertilization success rather than the binary acceptance-rejection response expected under self-incompatibility. This analysis excluded nearly 20% of donor-recipient combinations – a proportion that likely exceeds the frequency of incompatible crosses expected in our *B. rapa* experimental population. Consequently, this exclusion likely removed not only incompatible combinations, but also compatible pollen donors that performed poorly under pollen competition. Finally, unsuccessful pairs were not more closely related than successful ones, as would have been expected for a kin-recognition mechanism such as sporophytic self-incompatibility.

In animals, compatibility-based cryptic female choice is often interpreted as an adaptive mechanism of inbreeding avoidance, and in some cases outbreeding avoidance (inbreeding avoidance: Gasparini & Pilastro, 2011; Oliver & Evans, 2014; Fitzpatrick & Evans, 2014; Firman & Simmons, 2015; Rosengrave *et al*., 2016; outbreeding avoidance: Yeates *et al*., 2009; Gasparini *et al*., 2015). Under this scenario, females favor males that confer genetic fitness benefits relative to their own genotype. In our experiment, each plant represented a distinct genotype because individuals originated from distinct experimental populations. In plants, inbreeding avoidance has generally been attributed to self-incompatibility systems rather than to sexual selection among compatible partners (Guo *et al*., 2019; Genete *et al*., 2020). Although we found no evidence that cryptic choice mediated by pollen-pistil interactions in *B. rapa* reject either closely related pollen donors, or particularly distant ones, we cannot exclude the possibility that once the sporophytic self-incompatibility system has eliminated the most closely related mates, interaction between pollen tubes and pistil tissues enables paternity bias according to more subtle differences in relatedness. Testing this hypothesis will require quantifying pollen-pistil interaction scores and offspring fitness across controlled gradients of genetic relatedness between pollen donors and recipients. More broadly, our results suggest that compatibility-based cryptic female choice may be more diverse than initially envisioned, thereby opening new avenues for investigating post-pollination mate choice processes in plants.

### Style length exacerbates cryptic female choice

Style length emerged as a key pistil trait modulating the compatibility-based female choice process mediated by pollen tube length after interaction with pistil tissues. Under pollen competition, recipients with longer styles biased paternity more strongly toward pollen donors that produced longer pollen tubes after the interaction with pistil tissues in the absence of competition. Recently, it was shown in *B. rapa* that plants with short and long styles bias paternity toward donors with opposite *in vitro* pollen scores, including pollen tube growth rate, germination rate, and pollen size (Chenin *et al*., 2026). However, as mentioned above, these individual male scores do not predict pollen-pistil interaction traits (either alone or in interaction with female individual scores). Our results therefore complement previous findings, and reveal that style length is involved in at least two distinct forms of cryptic female choice, emphasizing the central role of style morphology in post-pollination sexual selection.

While style length acted here as a modulator of the strength of compatibility-based cryptic female choice, our assessment of pollen tube growth after interaction with pistil tissues likely reflects underlying cellular and/or molecular interactions between them. Indeed, pollen tube growth after interaction with pistil tissues was independent of the morphology of the pistil deposited onto the medium, suggesting that the observed patterns were not solely driven by morphological differences among pistils. Furthermore, our semi-*in vivo* protocol likely captured aspects of chemical pollen tube attraction toward unfertilized ovules, as greater distances between ovules and the tip of the excised style were associated with increased pollen tube growth within the medium. Although our understanding of cellular and molecular interactions between pollen and pistil tissues are increasing (Palanivelu & Tsukamoto, 2012; Higashiyama & Takeuchi, 2015; Johnson *et al*., 2019), the mechanisms underlying the compatibility-based cryptic female choice pattern reported here remain unknown. Variation in style length may influence the strength of compatibility-based female choice occurring under pollen competition because such phenotypic variation may be accompanied by quantitative or qualitative differences in female-emitted resources and molecular guidance cues necessary for pollen tube growth. Although our experimental design did not include comparisons of protein expression among styles differing in length, our finding that pollen tubes emerging closer to the center of the style were longer may indicate spatial heterogeneity within the style itself, consistent with previous molecular studies (Holdaway-Clarke & Hepler, 2003; Johnson *et al*., 2019).

Our study provides empirical evidence for a previously undescribed form of cryptic female choice during the post-pollination phase of plant reproduction underlined by physical contact between pollen tubes and pistil tissues. Paralleling post-mating sexual selection processes described in some animals, female preference varied according to pollen donor identity, consistent with a scenario of compatibility-based female choice, rather than with directional models of female choice. Together with recent empirical findings in *B. rapa* (Chenin *et al*., 2026), our results identify the post-pollination phase as a key arena for sexual selection in plants, offering multiple opportunities for cryptic female choice to occur. A critical next step will be to test whether the compatibility-based female choice reported here is adaptive, for example by contributing to inbreeding or outbreeding avoidance, and to quantify the relative contributions of both the previously described processes in *B. rapa* and the compatibility-based process to variation in female fitness. Furthermore, characterizing pistil molecular composition in relation to pistil morphology, combined with experimental manipulation of pistil structure in semi-*in vivo* assays, will be essential for understanding whether and how the fine-scale dialogue between pollen and pistil tissues underlies cryptic female choice in plants.

## Material and Methods

### Biological material

Our experiment was conducted using *Brassica rapa*, an annual, hermaphroditic, and generalist insect-pollinated angiosperm species. Plants originated from the Wisconsin Fast Plants™ lineage (WFP™, Carolina Biological Supply Company, Burlington, NC, USA; Williams & Hill, 1986), established from multiple wild cabbage populations collected across North America. This line underwent selection for several generations for its rapid life cycle (∼8 weeks) and high fecundity under greenhouse-grown conditions, while maintaining substantial genetic and phenotypic diversity (Williams & Hill, 1986). Like other members of the Brassicaceae, *B. rapa* possesses a sporophytic self-incompatibility system that prevents self-fertilization. Nevertheless, self-incompatibility is not complete with a low frequency of B. rapa plants able to self, including in the WFP™ line (Williams & Hill, 1986; Takayama & Isogai, 2005; Nasrallah, 2019; Ramos & Schiestl, 2019). Flowers typically exhibit four superior and two inferior stamens spatially separated from the pistil (herkogamy), thereby limiting self-pollination. These characteristics allowed precise control over the number of pollen donors deposited on the stigma during manual pollination, ensuring that pollen competition occurred primarily among outcrossed individuals. Moreover, the *B. rapa* WFP™ lineage has been successfully used in semi-*in vivo* assays (Lorbiecke, 2012), making it a suitable system to test whether pollen-pistil interactions can underlie cryptic female choice.

Plants used in our experiment originated from five experimental populations, established from WFP™ seeds purchased from the University of Wisconsin in 2019, that have evolved under strong polygamous conditions for three to four generations. At each generation, seeds from each experimental polygamous population were sown to generate a pool of approximately 85 parental plants (using the same growing conditions as described below for our experiment), which were used to establish the next generation through controlled manual pollination. Three flowers per parental plant were pollinated using a pollen mixture composed of a pollen mix collected across all 85 individuals and including twelve harvested stamens per plant. Seeds resulting from these pollinations (after three to four generations) constituted the five experimental “source populations”, from which both pollen donor and pollen recipient plants used in our experiment were derived.

### Experimental design

Several components of our experimental framework were described in Chenin *et al*. (2026). Data from *in vitro* pollen trait measurements, pistil trait measurements, and paternity analyses were shared between the two studies (Chenin *et al*., 2026). However, the semi-*in vivo* dataset was generated specifically for the present study to address original research questions. Mature pollen donor and pollen recipient plants were organized into “mating groups” to perform manual pollination. The experiment was conducted in January 2023 in the greenhouses of the CNRS experimental platform in Montpellier, France. Pollen donor and recipient plants were sown in three temporal blocks spaced one week apart to ensure a continuous supply of sexually mature plants throughout the experiment. Plants were grown individually in 0.7 L pots filled with a sterilized soil mixture under standardized greenhouse conditions, maintained in a water-saturated environment with continuous light at 25 °C. To prevent undesired cross-pollination, plants were enclosed in insect-proof mesh cages and spaced apart to avoid physical contact during manipulation. Plants were randomly selected on the evening prior to manual pollination and had to meet the following criteria to be included into a mating group: pollen recipient plants had to exhibit two virgin flowers about to open (hereafter “recipient flowers”), and pollen donor plants had to exhibit one flower with non-dehiscent stamens (“donor flower”). The five pollen donor plants of a given mating group originated from our five source populations and from the same temporal sowing block, as did the five recipient plants. These criteria were applied consistently across all mating groups to standardize our results by flower age, but donors and recipients were chosen randomly among plants that met these criteria. We established fourteen mating groups throughout the experiment, totaling 70 pollen donors and 70 pollen recipient plants.

Manual pollinations were performed on one or several mating groups per day, thus forming a “pollination session”. Two superior stamens from the donor flowers were collected early in the morning and placed in Petri dishes under a lamp for 3–4 h to accelerate stamen dehiscence. Pollen was then thoroughly homogenized under a binocular microscope using a brush and used to pollinate recipient flowers. During hand pollination, stigmas were saturated with the pollen mixture from the five pollen donor plants. Because the number of deposited pollen grains exceeded the number of ovules, pollen donors effectively competed for fertilization. The quantity of pollen deposited on stigmas was largely saturating and standardized among pollen recipient plants by pollinating the first recipient flower of each plant in a predetermined random order, and the second flower in the reverse order. The brush used for manual pollination was replaced and reloaded with pollen between the two pollination rounds to limit cross-contamination among pollen recipient plants. Despite these precautions, a significant proportion of paternities were attributed to pollen recipient plants within the mating groups due to pollen transfer between successive pollinations; seeds sired by pollen recipient plants within the mating groups were removed from subsequent analyses (see below). Although our pollination protocol was optimized to minimize selective processes during the pre-pollination phase, the exact number of pollen grains deposited from each pollen donor plant was not strictly controlled. Because variation in pollen production can substantially influence paternal fertilization success, one superior stamen was collected from each donor flower to estimate pollen production per donor plant (see below). Pollen production was controlled for in all subsequent statistical analyses.

### Assessing pollen-pistil interactions in semi-in vivo assays

Semi-*in vivo* assays aim at quantifying both pollen tube growth and ovule-targeting ability following interactions between pollen and pistil tissues. Specifically, the assay consists of pollinating freshly excised virgin stigmas and styles placed in a growing medium with fresh pollen, and subsequently observing the ability of pollen tubes to grow and target unfertilized excised ovules positioned around the base of the style after a standardized growth period. This experimental design was adapted from a landmark molecular biology study demonstrating that pistil tissues capacitate pollen tubes to locate ovules (Higashiyama *et al*., 1998; Palanivelu & Preuss, 2006; Stewman *et al*., 2010; Lorbiecke, 2012), by comparing pollen tubes’ ability to target the ovule micropyle after growing through pistil tissues versus pollen germinated exclusively *in vitro* (Higashiyama *et al*., 1998).

In our study, we adapted semi-*in vivo* assays to test whether molecular interactions between pollen and pistil tissues are involved in cryptic female choice. To this end, we followed experimental approaches classically applied to study cryptic female choice in externally fertilizing animals, based on fully factorial designs exploring all donor-recipient combinations (Urbach *et al*., 2005; Rosengrave *et al*., 2008, 2016; Butts *et al*., 2012; Evans *et al*., 2013; Oliver & Evans, 2014; Geßner *et al*., 2017; Poli *et al*., 2019). For each pollen donor-recipient plant combination within a mating group (*n* = 25), pollen collected from donor plants was deposited onto a single virgin pistil from each recipient plant in the semi-*in vivo* medium. Pollen tube growth and ovule-targeting ability were therefore assessed in the absence of direct competition between donors. Each pollen donor-recipient combination was represented once, resulting in a total of 350 semi-*in vivo* assays. Pistils from recipient plants and pollen from donor plants (*n* = 70) were collected on the same day as the pollination session. On each recipient plant, five pistils were sampled from freshly opened flowers that had been open for one to two days. On donor plants, pollen was collected from one superior stamen per donor flower. Although pistil collection was not strictly standardized by flower age, we prioritized the youngest flowers located at the top of the inflorescence to maximize the likelihood that pistils were virgin. Prior to pistil collection, stamens were carefully removed with tweezers to prevent accidental self-pollen deposition during sampling. Pistils were stored individually in 1.5 mL Eppendorf tubes, and semi-*in vivo* assays were conducted immediately after sampling. For one of the fourteen mating groups, pistils could not be collected from recipient plants, resulting in a final total of 325 pistils and stamens used for the assays.

The *in vitro* growth medium (PGM) on which excised pistils and unfertilized ovules were placed was adapted from Schreiber & Dresselhaus (2003) and optimized for *B. rapa* (Barbot *et al*., 2025; Chenin *et al*., 2026). To prepare 100 mL of PGM, we dissolved 10 g of sucrose and 3 g of PEG 6000 (polyethylene glycol) in 98 mL of distilled water, and then added 0.5 mL of 0.5% H₃BO₃, 0.5 mL of 0.1 M CaCl₂, 2.55 µL of 0.1 M KH₂PO₄, and 0.5 g of agar (0.5%). Agar concentration was adjusted to ensure medium stability during pistil manipulation and throughout the duration of the semi-*in vivo* assay. Pistils were excised at the style-ovary junction using a razor blade and placed horizontally on 200 µL of PGM deposited on a microscope slide. Ovaries were dissected along the carpellary junction under a binocular microscope, and eight ovules were carefully extracted by cutting the funiculus at its base using tweezers and an insect needle. Ovules were arranged in a semicircle around the excised style base at a distance of approximately 1.5 mm on the PGM (mean ± SD: 1.64 ± 0.30 mm; Figure 1). This spacing allows pollen tubes to detect and respond to female gametophyte guidance cues after capacitation by stigma and style tissues, as reported in several angiosperm species (Higashiyama *et al*., 1998; Palanivelu & Preuss, 2006; Stewman *et al*., 2010), including our *B. rapa* WFP™ lineage (Lorbiecke, 2012). Because ovule-style distance was not fully standardized and may influence pollen tube growth and ovule targeting ability, it was included as a random effect in subsequent analyses (see below). Ovules were spaced approximately 0.5–0.8 mm apart (mean ± SD: 0.67 ± 0.17 mm), with the micropyle oriented toward the excised style base. The four ovules positioned on the left side of the style were oriented to the right, whereas those on the right side were oriented to the left (Figure S1). After placement of pistils and ovules on the PGM, the entire stigma surface was pollinated using an insect needle for each donor-recipient combination within a mating group (*n* = 25). Pollen deposition was optimized to obtain 10–50 pollen tubes emerging from the excised style base, enabling individual tracking of pollen tubes after emergence from pistil tissues, and reliable quantification of pollen traits (see below). Because pollen from the focal stamen had previously been used for germination assays (see below), pollen quantity was insufficient to pollinate all five pistils in 20 of the 325 semi-*in vivo* assays. In these cases, stigmas were supplemented with pollen collected from another flower from the same inflorescence as the donor flower.

Microscope slides were incubated at 25°C in Petri dishes containing a 3.5 × 7 cm cotton pad moistened with 2 mL of water to maintain humidity and prevent desiccation of pistil tissues. Images were captured after 26–30 hours of incubation (mean ± SD: 28.18 ± 1.61 hours), once pollen tubes had emerged from the excised style base. Although incubation time was not fully standardized, statistical analyses revealed no effect of incubation duration on pollen tube growth or orientation (see below). This result is consistent with our observations that pollen germination and tube growth ceased after 24 hours, as confirmed by comparing images taken at 24 and 26 hours. Images were acquired using an optical microscope (Olympus CX43, Olympus Corporation, Tokyo, Japan) equipped with a 10× objective, allowing visualization of pistil tissues, pollen tubes emerging from the excised style base and growing in the medium, and the eight ovules. When all components could not be brought into focus simultaneously, multiple images were acquired using a 5× objective.

### Image processing and pollen-pistil interaction trait measurements for each donor-recipient pair

By processing images acquired from semi-*in vivo* assays (Figure 1A), we aimed to quantify variation among pollen donors in pollen tubes growth and ovule-targeting abilities to test whether pollen-pistil interactions mediate cryptic female choice. To this end, we quantified three quantitative traits designed to capture multiple components of pollen tube growth and orientation toward ovules after capacitation by pistil tissues (referred to as “pollen-pistil interaction traits”). For five randomly selected pollen tubes per image, we measured: (i) pollen tube length, defined as the total length from the emergence point at the excised style base to the tube tip; (ii) pollen tube straightness, calculated as the ratio between the straight-line distance separating these two points and the total pollen tube length; and (iii) the remaining distance to the nearest micropyle, defined as the distance between the pollen tube tip and the nearest micropyle (Figure 1B).

Images were analyzed using a semi-automated ImageJ macro developed for this study (version 2.14.0; Schneider *et al*., 2012). For each acquired image, the scale was first calibrated to convert pixel measurements to micrometers, accounting for differences in microscope magnification (10× or 5× objectives). The excised style base was manually delineated using one or multiple line segments, along which five random coordinates were generated to select pollen tubes for measurements of pollen-pistil interaction traits. For each random coordinate, the nearest pollen tube emerging from the excised style base was selected, and its emergence point was recorded. Because pollen tube growth on the medium was often non-linear, pollen tube length was estimated as the sum of successive linear segments tracing the trajectory from the emergence point to the growth tip (Figure 1B). Pollen tube straightness was calculated as the ratio of the Euclidean distance between these two points and the pollen tube length (Figure 1B). Remaining distance to the nearest micropyle was estimated as the Euclidean distance between the pollen tube tip to the nearest recorded micropyle (Figure 1B). Pollen tubes emerging from the excised style base were observed in 174 of the 325 analyzed images, yielding a total of 712 measured pollen tubes, with an average of 4.09 (SD ± 1.39) pollen tubes per pollen donor-recipient combination. Within each mating group, an average of 13.38 maternal-paternal combinations (SD ± 4.65) were measured out of the 25 expected combinations. The first three mating groups underrepresented due to initially lower success in performing technically demanding manual semi-*in vivo* pollinations. However, excluding these three mating groups from the statistical analyses did not qualitatively affect the results.

On each acquired image, we quantified three uncontrolled positionnal factors potentially influencing pollen tube growth and ovule-targeting ability: (i) variation in ovule positioning around the excised style base, calculated as the mean of the Euclidean distances between the eight micropyle coordinates and each pollen tube’s emergence point; (ii) the relative distance between the pollen tube emergence point and the center of the excised style base, defined as the Euclidean distance between these two points divided by half the delineated style base length; and (iii) the distance pollen tubes traversed through pistil tissues before emergence onto the medium, estimated as the sum of stigma and style lengths. These factors were included as covariates in our statistical analyses, and their effects on variation in pollen tube length, pollen tube straightness, and the remaining distance to the nearest micropyle were tested (see below).

### Pistil traits measurements on recipient plants

Protocol of pistil traits measurement was previously described in Chenin *et al*. (2026). Pistil trait measurements were conducted on two pistils specifically sampled for this purpose from each recipient plant, as well as on the five pistils collected for the semi-*in vivo* assays. The two additional pistils were sampled from virgin flowers on the day of each pollination session. Because pistil morphology changes with flower age, flower buds expected to open shortly were marked the day before sampling. This protocol standardized pistil trait measurements with respect to flower age while minimizing the risk of prior pollen deposition on the stigma. The five pistils used for the semi-*in vivo* assays were measured under an optical microscope (Olympus CX43; Olympus Corporation, Tokyo, Japan), whereas the two additional pistils were measured under a binocular microscope. These methodological differences, as well as variation in sampling dates among mating groups, were accounted for in our statistical analyses (see below). In total, we collected 465 pistils on 70 pollen recipient plants, and measured 6.64 (± 1.30) pistils on average per recipient plant (± SD).

Pistils were individually stored in 1.5-mL Eppendorf tubes containing a tissue fixation solution composed of ethanol, methanol, and acetic acid (EMA; 1:1:3). Images were acquired with pistils positioned longitudinally under a binocular microscope equipped with a camera. Stigmas were then carefully excised with a razor blade and imaged from an apical view. Pistil were humidified with EMA solution during imaging to prevent tissues deformation caused by desiccation. Pistil traits were measured by analyzing images using ImageJ software (Schneider *et al*., 2012). Longitudinal images were used to measure the lengths and widths of the three morphological components of the pistil: stigma length (from the apex to the stigma-style junction), style length (from the stigma-style junction to the style-ovary), and ovary length (from the style-ovary junction to the base of the ovary). Widths were measured at the midpoint of each structure. Stigma surface area was determined from apical images. Because stigma area, ovary length, and ovary width could not be measured from pistil used in the semi-*in vivo* assays, only two measurements per pollen recipient were available for these traits, whereas seven measurements per recipient were obtained for the remaining pistil traits.

### Pollen traits measurements on donor plants

Protocol of *in vitro* pollen traits measurement was previously described in Chenin *et al*. (2026). In brief, *in vitro* pollen germination assays and pollen trait measurements were conducted using pollen collected from one superior stamen harvested in the donor flower of each donor plant (*n* = 70). Pollen was extracted immediately after sampling and deposited onto 20 µL of PGM on a microscope slide. The composition of the medium differed from that used in the semi-*in vivo* assays, as agar was omitted from the preparation (see above). Slides were placed individually in Petri dishes containing a 3.5 × 7 cm paper strip moistened with 2 mL of distilled water to prevent desiccation and incubated under standardized conditions at 20 °C. Images were acquired after one, three, and five hours of incubation using an optical microscope with a 10× objective (Olympus CX43; Olympus Corporation, Tokyo, Japan). Five images were captured at different locations on each slide to increase the number of pollen grains measured while avoiding repeated measurements of the same pollen grains. Pollen traits were assessed by analyzing images with ImageJ software (Schneider *et al*., 2012). For each image, we assessed: (i) germination rate, by classifying and counting pollen grains as germinated or non-germinated; (ii) pollen size, by measuring the diameter of three germinated and three non-germinated pollen grains; and (iii) pollen tube growth rate, by measuring the pollen tube length of the three germinated grains used for diameter measurements and two additional germinated pollen grains. A pollen grain was considered germinated when the pollen tube reached at least half of the pollen diameter, following previously published pollen trait measurement protocols (Luza *et al*., 1987; Kakani *et al*., 2005). A small proportion of pollen grains were classified as sterile because they were substantially smaller and failed to produce a pollen tube. In most images, some pollen grains were classified as indeterminate due to aggregation or suboptimal focus, preventing accurate categorization.

Pollen traits were measured using a semi-automated protocol based on random coordinates generated for each image provided in Chenin *et al*. (2026). Across the three standardized time points, we measured on average (± SD) 28.31 ± 10.21 non-germinated pollen grains, 28.67 ± 10.78 germinated grains, and 46.24 ± 17.05 pollen tubes per pollen donor plant. One individual was excluded from statistical analyses because all pollen grains were classified as sterile, preventing trait measurement. As germinated and non-germinated pollen diameters were positively correlated (Spearman’s ρ = 0.57, p < 0.001), we report only results for the diameter of germinated pollen grains, hereafter referred to as “pollen size”. Germination rate was estimated by counting germinated (361.87 ± 218.54), non-germinated (482.26 ± 338.72), and sterile pollen grains (53.07 ± 107.02) per pollen donor plant on average (± SD). Although the protocol was optimized to minimize variation in pollen deposition among pollen donor plants, the exact number of pollen grains deposited on slides was not precisely controlled. As pollen density is known to influence pollen growth dynamics, the total number of pollen grains per image was included as a covariate in subsequent statistical analyses (see below). For pollen donor plants from the first three mating groups, images were captured using a 5× objective. Because measurement precision can vary with magnification, objective type was included as a random effect in subsequent analyses (see below).

### Pollen production on donor plants

Pollen production per pollen donor plant was estimated by counting pollen grains extracted from a single superior stamen from each donor flower (*n* = 70). Importantly, this stamen was distinct from those used for manual pollination, semi-*in vivo* assays, and pollen germination assays (see above). Pollen preparation for counting was adapted and optimized for *B. rapa* based on the protocol of Loublier *et al*. (1986). Stamens were collected on the evening of a pollination session before stamen dehiscence to prevent pollen loss. Stamens were then immediately stored individually in 1.5 mL Eppendorf tubes at – 80 °C. Pollen extraction protocol was fully described in Chenin *et al*. (2026). In brief, stamens tissues were first digesting using sulfuric acid and then crushed with a glass rod to release pollen from residual tissues. Subsequently, distilled water was added into samples, and repeated cycles of sonication and centrifugation were applied to disperse aggregated pollen grains (Chenin *et al*., 2026). Of the 70 stamens sampled, 66 were successfully processed and used to estimate pollen production per pollen donor plant.

Pollen production per pollen donor plant was quantified by counting pollen grains on Malassez slides (5 × 5 grid) under a 10× objective using an optical microscope (Olympus CX43; Olympus Corporation, Tokyo, Japan). For each sample, pollen was counted in two successive 10-µL aliquots of the pollen suspension deposited on the slide. Following standard cell-counting protocols, grains overlapping the left or bottom borders of a grid cell were excluded from the count. In the rare cases where pollen formed aggregates, the number of grains was visually estimated based on aggregate size (ranging from 10 to a maximum of 100 grains). A minority of sterile pollen grains were observed and excluded from pollen production estimates, as their lack of viability was confirmed by the absence of pollen tube emergence in germination assays (see above). In total, approximately 5,800 pollen grains were counted across the 66 collected stamens.

### Fruit collection, seed classification and germination

Dried fruits were collected four to five weeks after manual pollination, once the pollen recipient plants had completed their life cycle. Of the 140 hand-pollinated flowers, 112 fruits were collected and manually opened, and seeds were classified into four visually distinct categories: (i) viable seeds, with a smooth brown seed coat (86.8% of 2,510 seeds); (ii) aborted seeds, distinguishable from viable seeds by multiple depressions in the seed coat resulting from incomplete embryo development (1.0%); (iii) unfertilized seeds, with a transparent seed coat and no embryo (11.2%); and (iv) germinated seeds, with a small plantlet emerging (1.0%). The 2,205 collected viable and aborted seeds were germinated in Petri dishes containing 10 gL⁻¹ agar to obtain sufficient DNA material for genotyping from the resulting plantlets. Offspring from the same fruit were placed in the same Petri dish and spaced at least one centimeter apart to minimize density effects on seed germination. Seeds were incubated for seven days under controlled conditions (20–23 °C, 16:8 h photoperiod), yielding a germination rate of 97.3% among viable seeds. As all aborted seeds failed to develop into plantlets (*n* = 26), this category was considered consistent across samples. All resulting plantlets were subsequently genotyped to estimate the ovule fertilization success of each donor plant on each recipient plant.

### Genotyping parental and offspring tissues

Fresh leaves from pollen donor and recipient plants were collected for genotyping five to seven days after manual pollination. A detailed description of the genotyping protocol is provided in Barbot *et al*. (2025). Parental leaf samples were incubated (24 h at 35 °C) and 10–12 mg of resulted dried tissue was used for DNA extraction. All offspring obtained from seed germination were genotyped, representing a total of 2,122 individuals. Parental and offspring DNA were extracted using the NucleoMag Plant kit (Macherey-Nagel Inc., Allentown, USA). Biological samples were crushed with a 3 mm tungsten bead tissue homogenizer (Qiagen, Hilden, Germany) and incubated in a thermomixer containing lysis buffer and RNase. Following centrifugation, clarified lysates were transferred to deep-well plates (Thermo Fisher Scientific, Waltham, USA). DNA purification was performed using automated magnetic-bead extraction on a KingFisher® workstation (Thermo Fisher Scientific, Waltham, USA), followed by three sequential ethanol washes (35%, 35%, and 80%). Microsatellite genotyping was performed using a multiplex PCR with eight loci previously optimized for *B. rapa* and shown to be polymorphic within our experimental populations (Barbot *et al*., 2025). Amplifications were carried out using Applied Biosystems™ Master Mix (Thermo Fisher Scientific, Waltham, USA) on an Eppendorf Mastercycler Nexus GSX1 thermocycler applying the following program: an initial denaturation step of 15 min at 95 °C; 30 cycles of 30 s at 95 °C, 90 s at 58 °C, and 1 min at 72 °C, followed by a final extension of 30 min at 60 °C. PCR products were diluted 1:120 in distilled water, adding formamide and a size standard marker (GeneScan™ 500 LIZ™, Thermo Fisher Scientific, Waltham, USA). Capillary electrophoresis was conducted on an ABI 3500XL 24-capillary sequencer (Applied Biosystems®). Allele sizes were scored using GeneMapper® software (Thermo Fisher Scientific, Waltham, USA). Genotyping was repeated for 514 offspring and 26 parental samples due to low-quality allele calls resulting from amplification failures.

### Paternity analysis

Paternity analyses were conducted using CERVUS software (version 3.0.7; Marshall *et al*., 1998; Kalinowski *et al*., 2007). The program calculates the probability that a pollen donor plant was the true father of a given offspring based on the offspring’s genotype and the known maternal genotype within a mating group. For each offspring genotyped, a LOD (logarithm of the odds) score was calculated, corresponding to the ratio of the likelihood that a candidate pollen donor was the true father to the likelihood that he was not. Paternity analyses were performed independently for each mating group, and a relaxed criterion of 80% was applied to retain the most probable father. Offspring with missing data for at least three microsatellite markers were excluded from the analysis, corresponding to 203 paternities.

Criteria for paternity assignment were based on simulations of 1,000 hypothetical offspring genotypes from parents with known genotypes drawn from the allele frequencies of the population, allowing a 1% error rate estimated from observed mismatches between pollen recipient plants and their known offspring (mean mismatch = 0.524% across all eight markers). The mean non-exclusion probability across markers was 0.075, ranging from 0.049 to 0.141 across mating groups. Pollen recipient plants were included as potential fathers to account for possible pollen contamination. Across fourteen mating groups, 42.9% of the offspring were assigned to pollen recipient plants, ranging from 13.2% to 71.9%. Offspring assigned to pollen recipient plants were excluded from subsequent statistical analyses, corresponding to 723 attributed paternities. In total, 985 paternity assignments out of 2,122 genotyped offspring were retained for statistical analyses.

### Statistical Analysis

Estimates of individual and pair-specific interaction scores corrected for experimental heterogeneity Individual-level pollen traits (*in vitro* pollen traits and pollen production) were measured on one stamen from the donor flower, whereas individual-level pistil traits were measured on five flowers per individual. Pollen-pistil interaction traits specific to each donor-recipient pair are replicated on five pollen tubes for each semi-*in vivo* assay. In addition, all measures are affected by the specific conditions in which they took place (*e.g.*, sowing block, number of pollen grains deposited on the microscope slide, how far ovules were deposited from the excised style base in semi-*in vivo* assays). To obtain a measure characterizing the trait of an individual, or the interaction trait of a donor-recipient pair, we incorporated these factors in linear mixed-effects models (GLMMs), as well as an “individual” or “pair” random factor. The latter is the factor of interest, allowing us to characterize each individual or pair, based on all replicates, and removing variation arising from experimental conditions. Therefore, we extracted latent values of the “individual” or “pair” random effect from GLMMs and standardised them. They were referred to as “*in vitro* pollen and pistil scores” and “pollen-pistil interaction scores” in the text, and were used as predictors in individual-or pair-level models to characterize: (i) relationships between pair-specific interaction traits and *in vitro* pollen and pistil scores, and (ii) the relationship between pair-specific interaction scores and paternal fertilization success. All GLMMs were fitted using the `spaMM` package (version 4.6.39; Rousset & Ferdy, 2014) implemented in *R* (version 4.4.2; RCoreTeam, 2025). Below, we detail for each individual trait and score, the models from which we extracted the latent values.

#### Pair-specific pollen-pistil interaction scores

Our three pollen-pistil interaction traits were measured on five different pollen tubes for each donor-recipient pair. We modeled them using a Gamma error distribution (with log link), as they represent distances or ratios of distances bounded at zero. The model initially included linear fixed-effect covariates. Some were specific to each pollen tube: (i) the mean distance between the eight micropyles and the pollen-tube emergence point at the excised style base, and (ii) the relative distance between the pollen-tube emergence point and the center of the style base. Other covariates were shared among the five pollen tubes in the same image: (iii) the total distance traveled by the pollen tube through stigma and style tissues (*i.e.*, the sum of stigma and style lengths), and (iv) the time interval between the start of the semi-*in vivo* assay and image acquisition. For each interaction trait, we removed from the model the covariates with nonsignificant effect in order to avoid overfitting, leaving only (i) and (ii) in the three models. Random effects in the model included : semi-*in vivo* assay date, sowing block, population of origin of donor and recipient (in interaction), pollen donor identity, pollen recipient identity, and donor × recipient interaction. The donor × recipient term accounted for nearly all the variance jointly explained by the last four factors (Table 1). Therefore, we extracted the corresponding latent values from the models using the `ranef` function and used it as pair-specific interaction scores.

#### Individual *in vitro* pollen scores

Pollen traits *in vitro* (*i.e.*, germination rate, pollen size, and tube growth rate) were measured in one microscope slide per individual, where the pollen of one stamen from the donor flower was spread. Pollen size and square-root-transformed pollen tube growth rate were modeled assuming a Gaussian error distribution, whereas germination rate was expressed as a two-column response (number of germinated and non-germinated - including sterile - grains) and modeled using a binomial error. However, germination rates calculated including or excluding sterile pollen grains were highly and positively correlated (Spearman’s ρ = 0.97, p < 0.001), and subsequent analyses yielded consistent results regardless of whether sterile pollen was included. The GLMMs initially included fixed factors known to influence estimates of *in vitro* pollen traits (Holm, 1994; Pasonen & Käpylä, 1998; Boavida & McCormick, 2007): image acquisition time and pollen density on the slide – the latter as a quadratic term, as non-linear effects have been reported in the literature (Holm, 1994).

All fixed covariates were significant, and kept in the final model for germination rate and pollen size; whereas pollen density effects (linear and quadratic) were nonsignificant and removed from the pollen tube growth rate model. For all three individual *in vitro* pollen traits, random effects included sowing block, stamen sampling date, microscope objective, pollen measurer identity, source population, and pollen donor plant identity. The latent values for “donor plant identity” were extracted from the GLMMs and kept as individual-level estimates of *in vitro* pollen scores.

#### Individual pollen production scores

Pollen production was estimated based on pollen count on one stamen for each plant. The number of pollen grains counted per stamen was modelled using a Poisson distribution with log link, with no fixed-effect covariate and random effects: sowing block, stamen sampling date, source population, and pollen donor plant identity. The latent values for “donor plant identity” were extracted from the GLMMs and kept as an estimate of individual pollen production score.

#### Individual pistil scores

Recipient pistil traits (stigma area, style length, ovary length) were modeled as gaussian traits, with no fixed-effect covariates, and random effects included sowing block, pistil sampling date, microscope used for measurements, source population, and pollen recipient plant identity. The latent values for “recipient plant identity” were extracted from the GLMMs and kept as an estimate of individual pistil scores.

Relationship between pollen-pistil traits and individual in vitro pollen and pistil scores We tested whether pollen-pistil interaction traits depended on individual *in vitro* pollen and pistil scores estimated separately in donors and recipients respectively. To this end, we incorporated, among the linear predictors of pair-specific interaction traits (GLMM described above), donor pollen scores (pollen size, germination rate, pollen tube growth rate) and recipient pistil scores (stigma area, style length, ovary length) and their interactions (products), in addition to other covariates. The exclusion of all interactions between pollen and pistil scores did not significantly decrease the likelihood of the model, and similarly, none of the individual pollen scores or pistil scores had a significant effect (p-values were corrected for multiple testing using the false discovery rate method (FDR); *n* = 9 tests per pollen-pistil score model; Benjamini & Hochberg, 1995). In tables S1 and S2, we nevertheless report these nonsignificant effects in detail for the only interaction trait (pollen tube length after pollen-pistil interactions) that has an effect on paternities.

#### Multinomial models of fertilization success

In our experimental design, five pollen donors compete to fertilize seeds produced in each fruit. The numbers of seeds sired by each donor are a multinomial draw, constrained by the total number of seeds in the fruit. Each potential father sires on average 1/5 of this number, but siring success can be unequal among them. Our aim is to test whether such inequalities can be predicted based on interaction scores, *i.e*. covariates describing the performances of pollen donors when their pollen is put in contact with excised styles and ovules of the recipient, in semi-*in vivo* assays. To this end, we fitted a multinomial logit mixed-effect model implemented in the `pois4mlogit` function in the `spaMM` package (Rousset & Ferdy, 2014). This model predicts the number of seed sired by each donor, accounting for the influence of covariates both on its own fertilizing ability and on that of his competitors. It incorporates random effects to account for the same donor being represented on different recipients. Finally, when some predictors (interaction scores) are not available for a specific donor in a specific recipient, the relative success of the remaining donors on that recipient can still be used to inform the model without having to exclude the whole group.

We built a multinomial model of fertilization success including several linear predictors. The first linear predictor is a donor-specific score, pollen production. The trait was measured and corrected as described above, and then converted to a relative contribution score, dividing by the sum of all five males present in the pollen mix (when pollen production was not available for one donor, relative contribution scores were computed among the available ones). The other three linear predictors were the three pair-specific interaction scores (pollen tube length, pollen tube straightness, and remaining distance to the nearest micropyle). Remember that each score characterizes a donor-recipient pair and has been measured in semi-*in vivo* assays, and then corrected for experimental variation using GLMMs (see above). Random effects included pollen donor identity and the source populations of donor and recipient plants. The effect of each linear predictor was tested by sequentially removing each and performing LRTs (Table 1). Of these predictors, only one (pollen tube length after pollen-pistil interactions) turned out to have a significant (and positive) effect (see Results).

We then built an additional multinomial model to test whether the effect of the interaction scores could be modulated by pistil morphology. We constructed a model with donor-specific relative pollen production score, pair-specific interactions scores, and interactions between the latter and three recipient-specific pistil scores as predictors. It is important to realize that by construction, in this model, recipient-specific traits cannot appear as main effects, as the model predicts the *relative* success of five donors in competition on a common recipient – each will sire on average 1/5 of total seeds, irrespective of recipient scores. However, recipient scores may alter the relationship between interaction scores and paternities within a group of five competing donors, as represented by the interactions in our model. Interaction effects were tested using LRTs, with p-values corrected for multiple testing using the FDR method (*n* = 21 tests; Benjamini & Hochberg, 1995). Results from this complete model were consistent with those obtained from testing interactions individually in separate models, and pistil scores were not correlated (Figure 2). Consequently, we report only results obtained from the complete model in the main text (Table 4).

#### Effect of donor-recipient genetic relatedness on relative fertilization success

Genetic relatedness was estimated using SPAGeDi software (version 1.5; Hardy & Vekemans, 2002), based on eight microsatellite markers known to be variable in our experimental populations and also used for paternity assignment. To test the effect of pollen donor-recipient genetic relatedness on relative paternity success, we first computed six measures of relatedness between mates and retained the one with the lowest variance among both within-population and among-population mate pairs. By this logic, we retained Loiselle’s kinship estimator (Loiselle *et al*., 1995), which is indeed known to have low variance when population structure is weak (Vekemans & Hardy, 2004; Watts *et al*., 2007), as was the case here. Loiselle’s kinship estimator was included as a fixed effect in a multinomial model of relative fertilization success (described above), along with our three pollen-pistil interaction scores (pollen tube length, remaining distance to the nearest micropyle, and pollen tube straightness) and the relative pollen production of pollen donors. The effect of donor-recipient genetic relatedness on paternity success was tested using LRT comparing the full model with a reduced model excluding the Loiselle’s kinship estimator term. An effect of donor-recipient genetic relatedness might also be expected when donor and recipient belong to the same experimental population, but there is no strong within-population inbreeding (average within-population mate pair relatedness = 0.026).

## Aknowledgments

We are very grateful to Pascal Cosette, Jean-Claude Mollet, and Arnaud Lehner for their advice on conducting semi-*in vivo* assays in the early stages of this work. We thank Denis Orcel for plant care and Lena Galera for technical assistance with the semi-*in vivo* assays. We are grateful to Elsa Noël, Julia Centanni, and Armelle Kempf for their work on pollen and pistil trait measurements, and to Fleur Hamoir for assistance in genotyping. We acknowledge the ’PollenExtra’ platform of the ISEM lab (Montpellier, France) for providing access to microscopy equipment, and the ’Plateforme des Terrains d’Expériences du LabEx CeMEB’ (Montpellier, France) for logistical and technical support, including hosting the plants analysed in this study. This work was funded by an ERC Tremplin grant from the University of Montpellier to JT and by an ANR grant (ANR-23-ERCS-0010-01) to JT.

During the preparation of this work, the authors used ChatGPT in order to assist with rephrasing and improving the clarity of specific sentences. After using this tool or service, the authors reviewed and edited the content as needed and takes full responsibility for the content of the publication.

## Author contributions

Conceptualization, J.T., methodology, J.T., E.B., T.C., F.R. and P.D.; formal analysis, T.C., F.R., P.D. and J.T.; investigation, T.C., E.B., J.T. and A.M.; writing – original draft, T.C. and J.T.; writing – review & editing, T.C., J.T., P.D., F.R.; supervision, J.T.; funding acquisition, J.T.

## Data availability

The original datasets and the code used for statistical analyses are deposited to the Zenodo repository https://doi.org/10.5281/zenodo.21757983 and are publicly available as of the date of publication. Any additional information required to reanalyze the data reported in this paper is available from the lead contact upon request.

## Supporting Information

Figures S1–S2, Tables S1–S4, and Supporting references

