## Supporting Information for "Physical interactions between pollen and pistil tissues mediate cryptic female choice in *Brassica rapa*"

The following Supporting Information is available for this article: Figures S1–S2, Tables S1–S4, and Supporting references

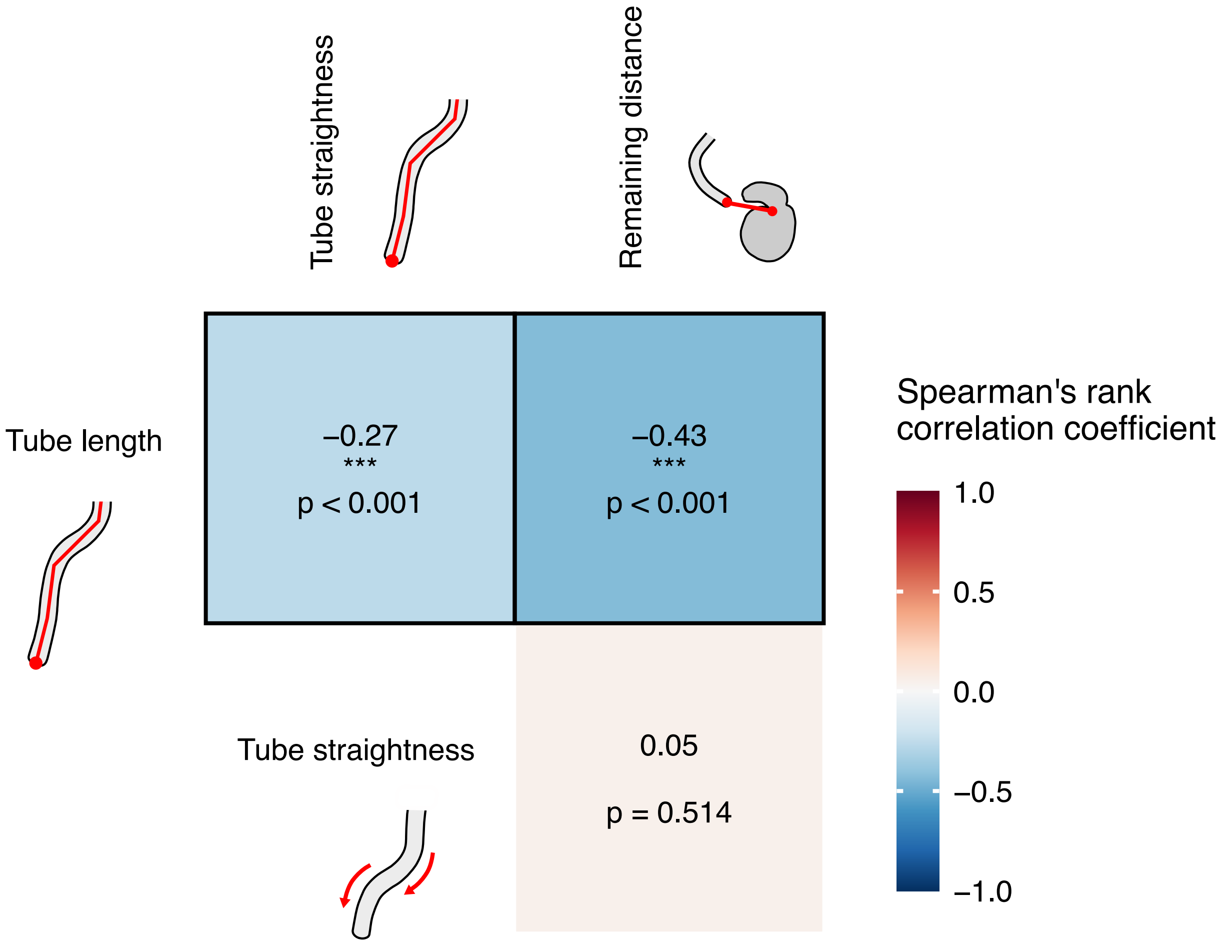


**Figure S1. Heatmap of correlations among pollen-pistil interaction scores**. Pollen-pistil interaction scores were standardized and correspond to predicted values extracted from the interaction between pollen donor and recipient identities, included as a random effect in GLMMs that corrected for experimental sources of variation (see Materials and Methods). Pairwise correlations were estimated using Spearman's rank correlation method. Color intensity reflects the strength and direction of correlations, with red and blue colors indicating positive and negative correlation values, respectively. *P*-values were adjusted for multiple testing using the FDR method ($n$ = 3 tests; Benjamini & Hochberg, 1995). Significant correlations are outlined in black. Asterisks indicate statistically significant p-values: * p < 0.05, ** p < 0.01, *** p < 0.001.

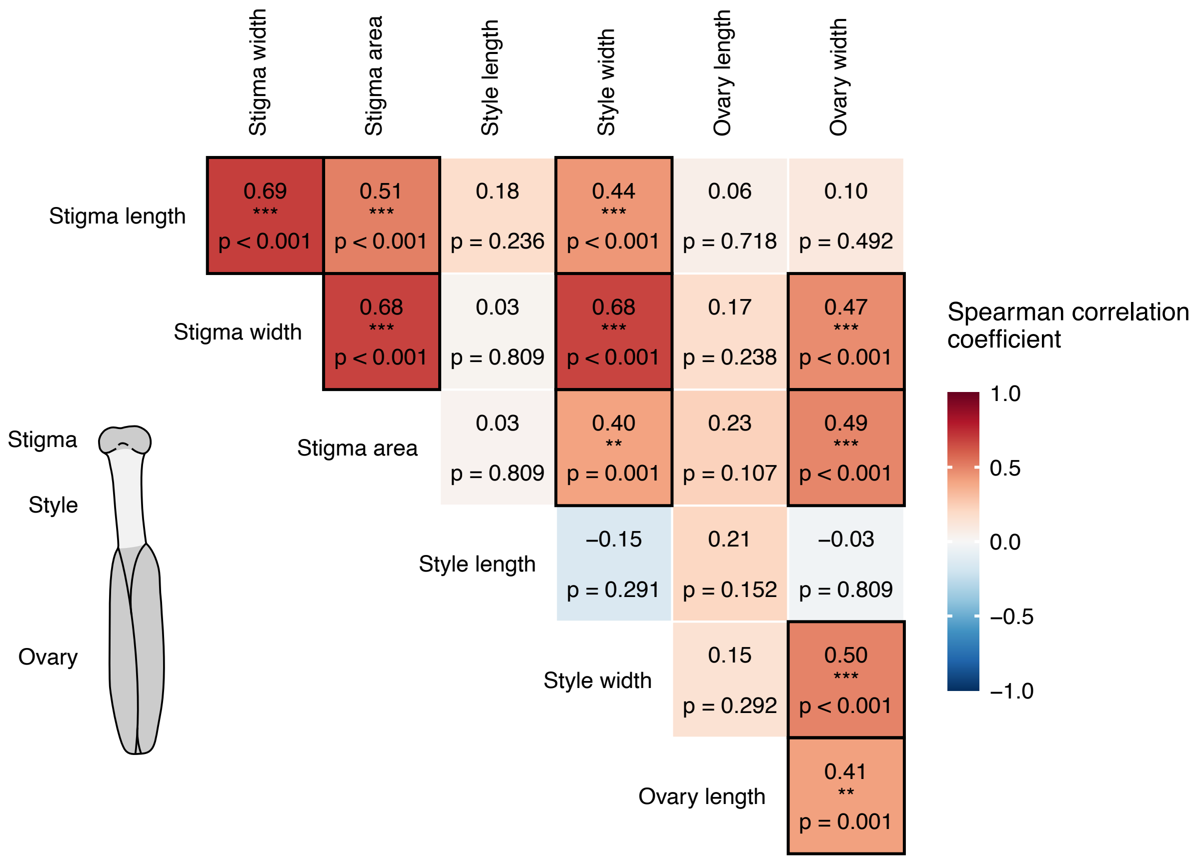


**Figure S2. Heatmap of correlations among pistil scores.** Pistil scores were standardized and correspond to predicted values extracted from pollen recipient identity included as a random effect in GLMMs that corrected for experimental sources of variation (see Materials and Methods). Pairwise correlations were estimated using Spearman's rank correlation method. Color intensity reflects the strength and direction of correlations, with red and blue colors indicating positive and negative correlation values, respectively. *P*-values were adjusted for multiple testing using the FDR method ($n$ = 21 tests; Benjamini & Hochberg, 1995). Significant correlations are outlined in black. Asterisks indicate statistically significant p-values: * $p < 0.05$, ** $p < 0.01$, *** $p < 0.001$.

| *In vitro* pollen and pistil scores | β ± SE | $\chi^2$ | p-value (LRT) |
|---|---|---|---|
| Pollen tube growth rate | −0.014 ± 0.044 | 0.093 | 0.760 |
| Pollen size | 0.061 ± 0.036 | 2.699 | 0.100 |
| Pollen germination rate | 0.046 ± 0.051 | 0.778 | 0.378 |
| Stigma area | −0.028 ± 0.038 | 0.545 | 0.460 |
| Style length | −0.027 ± 0.038 | 0.506 | 0.478 |
| Ovary length | 0.008 ± 0.038 | 0.039 | 0.844 |

**Table S1. Estimated effects of in vitro pollen and pistil scores on pollen tube length.** Pollen tube length was measured for each pollen donor-recipient (*n* = 325) combination in semi-in vivo assays after pollen–pistil interactions and analyzed as a response variable in a GLMM correcting for experimental sources of variation (see Materials and Methods). The effects of *in vitro* pollen and pistil scores on variation in pollen tube length were estimated in two separate models. The first model included pollen tube growth rate, pollen size, and pollen germination rate scores estimated for pollen donors as fixed effects (*n* = 70). The second model included uncorrelated components of pistil morphology as fixed effects: stigma area, style length, and ovary length scores estimated for recipient plants (*n* = 70). Pollen and pistil scores correspond to standardized predicted values extracted from the pollen donor or recipient identity term included as a random effect in GLMMs correcting for experimental sources of variation (see Materials and Methods). The significance of each in vitro pollen and pistil score was tested using LRTs comparing models with and without the effect. β values represent estimated effect sizes ± standard errors. Degrees of freedom were equal to one for all comparisons.

| *In vitro* pollen score × Pistil score | β ± SE | $\chi^2$ | p-value (LRT) | FDR p-value |
|---|---|---|---|---|
| Pollen tube growth rate × Stigma area | 0.054 ± 0.050 | 1.177 | 0.278 | 0.626 |
| Pollen tube growth rate × Style length | 0.023 ± 0.048 | 0.228 | 0.633 | 0.851 |
| Pollen tube growth rate × Ovary length | −0.018 ± 0.049 | 0.130 | 0.719 | 0.851 |
| Pollen germination rate × Stigma area | −0.027 ± 0.043 | 0.379 | 0.538 | 0.851 |
| Pollen germination rate × Style length | −0.076 ± 0.054 | 1.924 | 0.165 | 0.626 |
| Pollen germination rate × Ovary length | 0.092 ± 0.052 | 2.867 | 0.090 | 0.626 |
| Pollen size × Stigma area | 0.045 ± 0.037 | 1.454 | 0.228 | 0.626 |
| Pollen size × Style length | −0.007 ± 0.037 | 0.035 | 0.851 | 0.851 |
| Pollen size × Ovary length | 0.010 ± 0.039 | 0.059 | 0.808 | 0.851 |

**Table S2. Estimated effects of interactions between *in vitro* pollen and pistil scores on pollen tube length.** Pollen tube length was measured for each pollen donor-recipient combination (*n* = 325) in semi-in vivo assays after pollen–pistil interactions and analyzed as a response variable in a GLMM correcting for experimental sources of variation (see Materials and Methods). Pollen and pistil scores were estimated for each pollen donor (*n* = 70) and recipient plants (*n* = 70) respectively. The model included pollen tube growth rate, pollen germination rate, pollen size, stigma area, style length, and ovary length scores as fixed effects, along with all pairwise interactions between pollen and pistil scores (*n* = 9 interaction terms). Pollen and pistil scores correspond to standardized predicted values extracted from pollen donor or recipient identity, respectively, included as a random effect in GLMMs correcting for experimental sources of variation (see Materials and Methods). The significance of each interaction was tested using LRTs comparing models with and without the effect. β values represent estimated effect sizes ± standard error. Degrees of freedom were equal to one for all comparisons. *P*-values were corrected for multiple testing using the FDR method (Benjamini & Hochberg, 1995), based on *p*-values from all models performed (*n* = 9 tests).

| Predictor | β ± SE | $\chi^2$ | p-value (LRT) |
|---|---|---|---|
| Pollen tube length score | 0.155 ± 0.071 | 4.807 | 0.028* |
| Remaining distance score | −0.076 ± 0.067 | 1.269 | 0.260 |
| Pollen tube straightness score | −0.158 ± 0.088 | 3.206 | 0.073 |
| Relative pollen production score | 0.480 ± 0.578 | 0.661 | 0.416 |

**Table S3. Estimated effects of pollen-pistil interaction scores and relative pollen production score on paternal fertilization success after excluding donor-recipient pairs with less than 5% of fertilized seeds.** Pollen donor-recipient plant pairs with less thant five percent of fertilized seeds were excluded from the statistical analyses (see Materials and Methods). The model included the three pollen-pistil interaction scores as fixed effects, and the relative pollen production score of pollen donors as a covariate (see Materials and Methods). All variables were standardized before inclusion in the model. The significance of each pollen trait was tested using LRTs comparing models with and without the effect. β values represent estimated effect sizes ± standard error. Degrees of freedom were equal to one for all model comparisons. Asterisks indicate statistically significant p-values: * $p < 0.05$, ** $p < 0.01$, *** $p < 0.001$.

| Pollen tube length score × Pistil scores | β ± SE | $\chi^2$ | p-value (LRT) |
|---|---|---|---|
| Pollen tube length × Stigma area | 0.148 ± 0.115 | 1.671 | 0.196 |
| Pollen tube length × Style length | 0.205 ± 0.100 | 4.364 | 0.037* |
| Pollen tube length × Ovary length | 0.036 ± 0.084 | 0.181 | 0.671 |

**Table S4. Estimated effects of the interactions between pollen tube length score and stigma area, style length and ovary length scores, excluding donor-recipient pairs with less than 5% of fertilized seeds**. Pollen donor-recipient plant pairs with less than five percent of fertilized seeds were excluded from statistical analyses (see Materials and Methods). The model included pollen tube length score as a fixed effect, along with its interaction with stigma area, style length, and ovary length scores, three uncorrelated components of pistil morphology (see Materials and Methods). Relative pollen production score estimated for each pollen donor was included as a covariate in the model (see Materials and Methods). Pollen tube length interaction score was obtained for each pollen donor-recipient plant combination (semi-*in vivo* assays, $n = 325$), and pistil scores were obtained from several flowers of the recipient plant ($n = 70$). All variables were standardized before inclusion in the model. The significance of each interaction between pistil scores and pollen-pistil interaction scores was tested by comparing models with and without the effect using LRTs. β values represent estimated effect sizes ± standard error. Degrees of freedom were equal to one for all comparisons. Asterisks indicate statistically significant p-values: * $p < 0.05$, ** $p < 0.01$, *** $p < 0.001$.
